# The Potential Regulation of A-to-I RNA editing on Genes in Parkinson’s Disease

**DOI:** 10.1101/2022.09.29.510217

**Authors:** Sijia Wu, Qiuping Xue, Xinyu Qin, Liyu Huang

## Abstract

Parkinson’s disease (PD), the second most common neurodegenerative disorder, was characterized by alpha-synuclein pathology and dopaminergic neuron degeneration. In previous studies, multiple genes have been demonstrated to involve in the regulations of these two processes, including *EIF2AK2*, *AGO2*, *MCL1*, *EEF1A1*, and *AIMP2*. The molecular mechanisms to mediate the transcript diversity of these genetic biomarkers were important to understand neurodegenerative pathogenesis and helpful for treatment design. In this study, we analyzed 372 PD patients to identify 9,897 A-to-I RNA editing events probably responsible for the controls of 6,286 genes. Due to the most potentially trans-regulatory associations between RNA editing events and genes, we tried to explain one possible pathway from the view of disturbed miRNA regulations on genes due to A-to-I RNA editing events. Specifically, we identified 72 RNA editing events probably interfering in miRNA regulations on their host genes, eight RNA editing events possibly altering miRNA competitions between their host genes and 1,146 other genes, and one RNA editing event modifying miRNA seed regions to potentially disturb its regulations on four genes. All the analyses revealed 25 RNA editing biomarkers in Parkinson’s pathogenesis through probably interfering in miRNA degradations on 133 PD-related genes.

## INTRODUCTION

Parkinson’s disease (PD), characterized by motor symptoms, olfactory disorder, somnipathy, emotional problems, and so on [1–2], is the second most common neurodegenerative disorder [3–4]. Its pathogenesis was verified to be modulated by genetic factors through key cellular processes [5–7], except for age and environmental exposures [8–9]. The first identified genetic factor was the alpha-synuclein gene [10], whose mutations were associated with PD risk [11]. Involved in the expression control of this gene, argonaute 2 (*AGO2*) was dys-regulated in PD patients [12]. And the inflammation-associated serine-threonine kinase (*EIF2AK2*) has been demonstrated to link with the phosphorylation and toxicity of alpha-synuclein [13]. Beside the alpha-synuclein pathology, the selective degeneration of dopaminergic neurons was also considered as a pathological manifestation of Parkinson’s disease. Some genes have been studied to be involved in this process [14], such as *MCL1* [15], *EEF1A1* [16], and *AIMP2* [17]. The studies of all these genetic biomarkers described their important functional roles in Parkinson’s disease. For these genes, it is necessary to understand their molecular mechanisms to reveal the underlying PD-related regulation pathways. Due to the increasing evidence of A-to-I RNA editing in neurological and neurodegenerative disorders [18–20] and its diverse roles in transcript diversity [21–22], we were inspired to explore the potential mechanisms of RNA editing effects on the expressions of genes, especially these PD-related genes.

For this goal, we collected 372 PD patients with RNA sequencing whole blood samples from a large consortium, Parkinson’s Progression Markers Initiative (PPMI) [23–24]. Based on these samples, we implemented the bioinformatics pipeline as shown in **Figure 1**, to identify the potential expression regulators, A-to-I RNA editing biomarkers, in Parkinson’s disease. Specifically, the first step was to identify the associations between genes and RNA editing events by two analysis methods. Next, we explored the mechanisms underlying these associations from the view of miRNA mediations, considering the possible involvements of RNA editing in miRNA dysregulations [21–22] and the gene degradations by miRNAs [25]. The analyses of the potential regulation processes were divided into three scenarios according to their locations, such as RNA editing events in miRNA targets to be associated with their host genes, RNA editing events possibly interfering in miRNA regulations on the competing genes, and RNA editing events in miRNA seed regions probably affecting miRNA-targeted genes. Form these analyses, we could discover specific RNA editing pathogenic biomarkers, and uncover their possible roles in Parkinson’s disease through the alterations of key PD-related genes.

**Figure 1.**
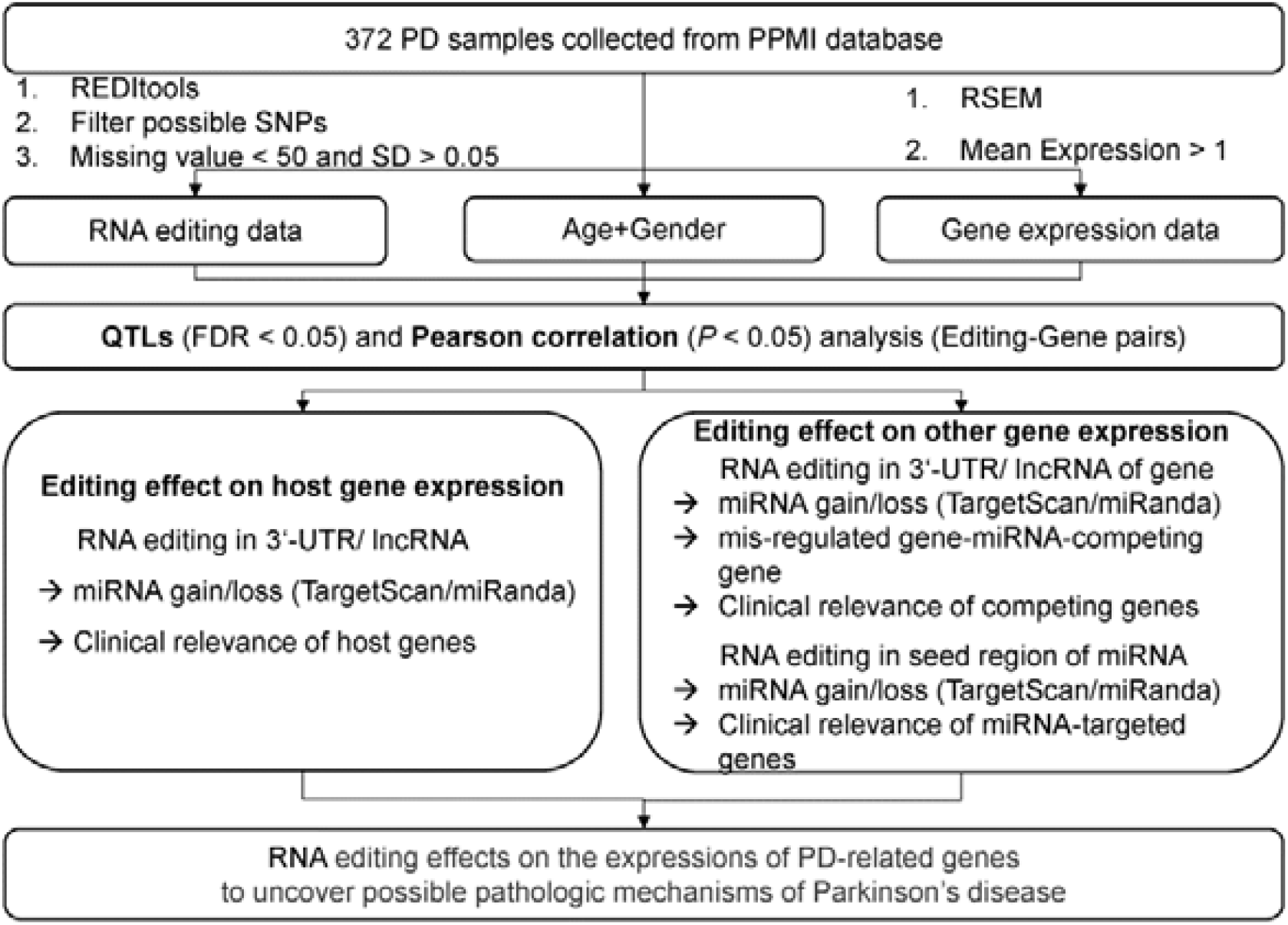
The flowchart of this study.

## MATERIALS AND METHODS

### Samples involved in this study

In total, we collected 372 PD patients with RNA sequencing blood samples from PPMI for the study of RNA editing effects on gene expressions in Parkinson’s disease. Moreover, there were another 169 healthy controls (HC) also downloaded from this consortium for the identification of PD specific biomarkers by their comparisons between healthy and PD groups. Beside the RNA sequencing data, the demographic information and clinical data (**Table S1**) of each individual were also included in this study for the clinical relevance of RNA editing and gene expression biomarkers. They included H&Y stages [26], alpha-synuclein observations, concentrations of amyloid β-protein [27], tau protein levels [28], and so on.

### A-to-I RNA editing detection

Based on the RNA sequencing samples, we first used STAR (v2.7.9a) [29] to align the reads to hg38 (GENCODE v36) reference files. Then we identified RNA editing events by REDItools [30] with default settings (e.g., minimal read coverage, 10; minimal quality score, 30; and minimal mapping quality score, 255). To ensure their confident identifications, we focused on known editing sites from REDIportal (December 2020) [31], removed possible SNP data from dbSNP151 [32], and filtered out the candidates with supporting reads under three or editing frequencies less than 0.1. Eventually, we selected A-to-I RNA editing events whose missing values were less than 50 and standard deviations were more than 0.05 as informative RNA editing events for further analysis. For all these informative RNA editing events, we analyzed their distributions in different types of genes, regions, and repeats by ANNOVAR [33], to reveal the reliability of our RNA editing identification pipeline.

### Gene expression quantification

With the aligned sequencing reads in bam files, we also performed quantitative analysis of transcriptome (TPM) using RSEM software [34]. The low abundant genes with mean expressions less than one were not considered in the following analysis, reserving 6,288 informative genes. Moreover, to reveal the potential effects of RNA editing on genes involved in Parkinson’s pathogenesis, we focused on 2,303 PD-related genes (**Table S2**) collected from DisGeNET [35], MalaCards [36], phenopedia [37], KEGG database [38], GWAS catalog [39], and one previous study [13].

### Correlation analysis between A-to-I RNA editing events and genes

To study the associations between A-to-I RNA editing events and genes, we conducted two kinds of analyses sequentially on them for reliability. They were Quantitative Traits Loci (QTLs) analysis by MatrixeQTL [40] using the criteria of *FDR* < 0.05 and Pearson correlation analysis with *P* < 0.05 for the significant associations. Specifically for QTL analysis, the factors of age and sex were considered as covariates, to improve its identification sensitivity. After analysis of the distributions between significant RNA editing events and genes, we discovered 1,203 RNA editing events in the 3’-UTRs of pre-mRNAs or lncRNAs to be associated with their host genes or miRNA competing genes, together with one RNA editing event locating in the miRNA seed regions to possibly affect the degradations of miRNA targets. They may disturb miRNA regulations on the expressions of genes, especially the PD-related genes.

### Analysis of potential miRNA regulation mechanisms involving in the RNA editing effects on genes

RNA editing events in miRNA binding targets, such as 3’-UTRs of pre-mRNAs or lncRNAs, may disturb miRNA regulations on their host genes. To study this, we first detected miRNA binding regions in wild-type and RNA-edited sequences by TargetScan (v7.2) [41] and miRanda (v3.3) [42]. The gain of miRNA binding targets was defined as the miRNA-RNA interactions occurring in the RNA-edited sequences but not in the wild-type sequences supported by both tools, and vice versa for the loss of miRNA binding targets. The effects of these RNA editing events on their host genes by disturbing miRNA regulations were further verified with the detected significant associations between editing frequencies and gene expressions. Later, for the further functions of the A-to-I RNA editing biomarkers through their potential effects on important genes, we identified the RNAs interacting with these genes from StarBase [43], and studied their enriched pathways by Metascape (v3.5) [44], DAVID(v6.9) [45] and Enrichr [46].

Moreover, RNA editing events in miRNA binding targets may not only introduce the changes in the expressions of their host genes, but also cause the alterations of other genes, due to the miRNA competitions of genes. To investigate this, we also used TargetScan and miRanda to identify miRNA targets in genes competing with the edited genes. The potential effects of A-to-I RNA editing events on these miRNA-competing genes were further verified by the detected significant associations between editing frequencies and the expressions of both genes. The further functional annotations of this kind of RNA editing events were analyzed similar as the above one.

Furthermore, some RNA editing events located in the seed regions (2-8nt) of miRNAs. They may alter their binding targets and degradation functions on specific genes, especially the collected PD-related genes. For this kind of RNA editing events, we also adopted similar procedures as above for their functional annotations.

## RESULTS

### A-to-I RNA editing events were involved in Parkinson’s disease through their effects on gene expressions

Our RNA editing detection pipeline first identified 25,799 informative RNA editing events. The majority of these editing events occurred in protein coding genes (93.80%), non-coding regions (99.77%), and Alu repeats (91.76%), as shown in **Figure 2A**. Their distributions were consistent with the preferred locations of RNA editing events reported in previous studies [21,47]. It partially revealed the reliability of the identified RNA editing events.

**Figure 2.**
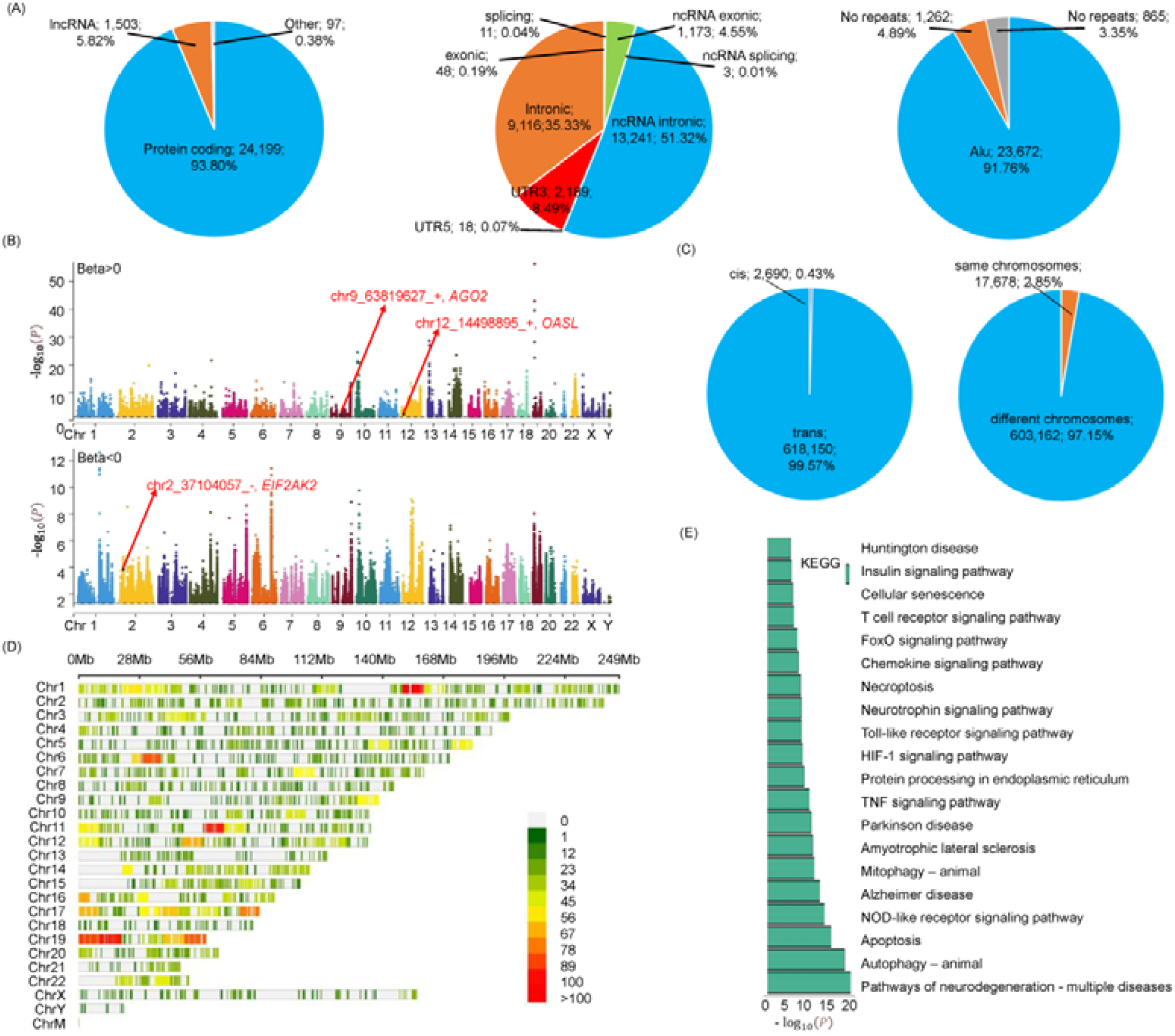
The overview of RNA editing effects on gene expressions. (A) The distributions of informative RNA editing events in genes, regions, and repeats. (B) Manhattan plots showing the effects of 9,897 RNA editing events on 6,286 genes by QTL (FDR<0.05) and Pearson correlation analyses (P<0.05). The top and bottom panels showed 137,563 positive and 483,277 negative associations respectively. (C) The distributions of these associations. (D) The density distributions of genes significantly associated with RNA editing events on chromosomes. They were highly dense in the regions of chromosomes 11 (chr11:60M-70M) and 19 (chr19:10M-20M). (E) The DAVID enrichment results of the PD-related genes probably affected by RNA editing events.

To explore the possible correlations between RNA editing events and genes, we performed both of the QTL (*FDR* < 0.05) and Pearson correlation (*P* < 0.05) analysis. In total, we identified 9,897 RNA editing events associated with 6,286 genes (**Table S3-5**) as the Manhattan plots shown in **Figure 2B**. These associations revealed the potentially preferred trans and cross-chromosomal RNA editing regulations on genes (**Figure 2C**). Moreover, for the genes associated with RNA editing events, we analyzed their distributions in genomic regions. It discovered the high density (Density > 100) in the regions of chromosomes 11 (chr11:60M-70M) and 19 (chr19:10M-20M), as shown in **Figure 2D**, different from the editing density in **Figure S1**. The potential mechanisms underlying these associations remained to be a problem needed to be addressed. Thus, in this study, we tried to explain that from the view of disturbed miRNA regulations on corresponding genes due to RNA editing events.

Of all these genes associated with A-to-I RNA editing, 820 genes were proposed previously to be linked with Parkinson’s disease. They may involve in the pathogenesis of this disease by the potential regulations of RNA editing biomarkers. To study the further functions, we performed enrichment analysis on these PD-related genes by DAVID (**Figure 2E, Table S6**). The enriched neurodegenerative diseases described the reliability of the collected genes related to Parkinson’s disease. On the other hand, the neurodegeneration related pathways that these genes were enriched in revealed the biological functions of these RNA editing events through their effects on the expressions of key genes.

From the analyses above, we could introduce one potential mechanism of RNA editing biomarkers in Parkinson’s pathogenies through their involvements in miRNA regulations on the key PD-related genes. Based on the locations of RNA editing events and their associated genes, the analysis was divided into three parts as shown in following sections one by one. They described the effects of RNA editing events in miRNA binding targets to be associated with their host genes or miRNA competing genes, and RNA editing events in miRNA seed regions to probably affect their downstream targets.

### A-to-I RNA editing events may affect their host genes by interfering in miRNA regulations

After the analysis about the locations of RNA editing events and corresponding genes, we identified 581 RNA editing events to be associated with their host genes. The underlying mechanisms could be explained from the view of altered miRNA regulations on their host genes due to the RNA editing events. To do this, we recognized 72 RNA editing events which were predicted to create 11,370 new miRNA binding targets and eliminate 815 original ones. Their effects on genes were verified by both of QTL and Pearson correlation analyses (**Table S7**), including *EIF2AK2* probably affected by an RNA editing event in the 3’-UTR of this gene, as shown in **Figure 3A-B**.

**Figure 3.**
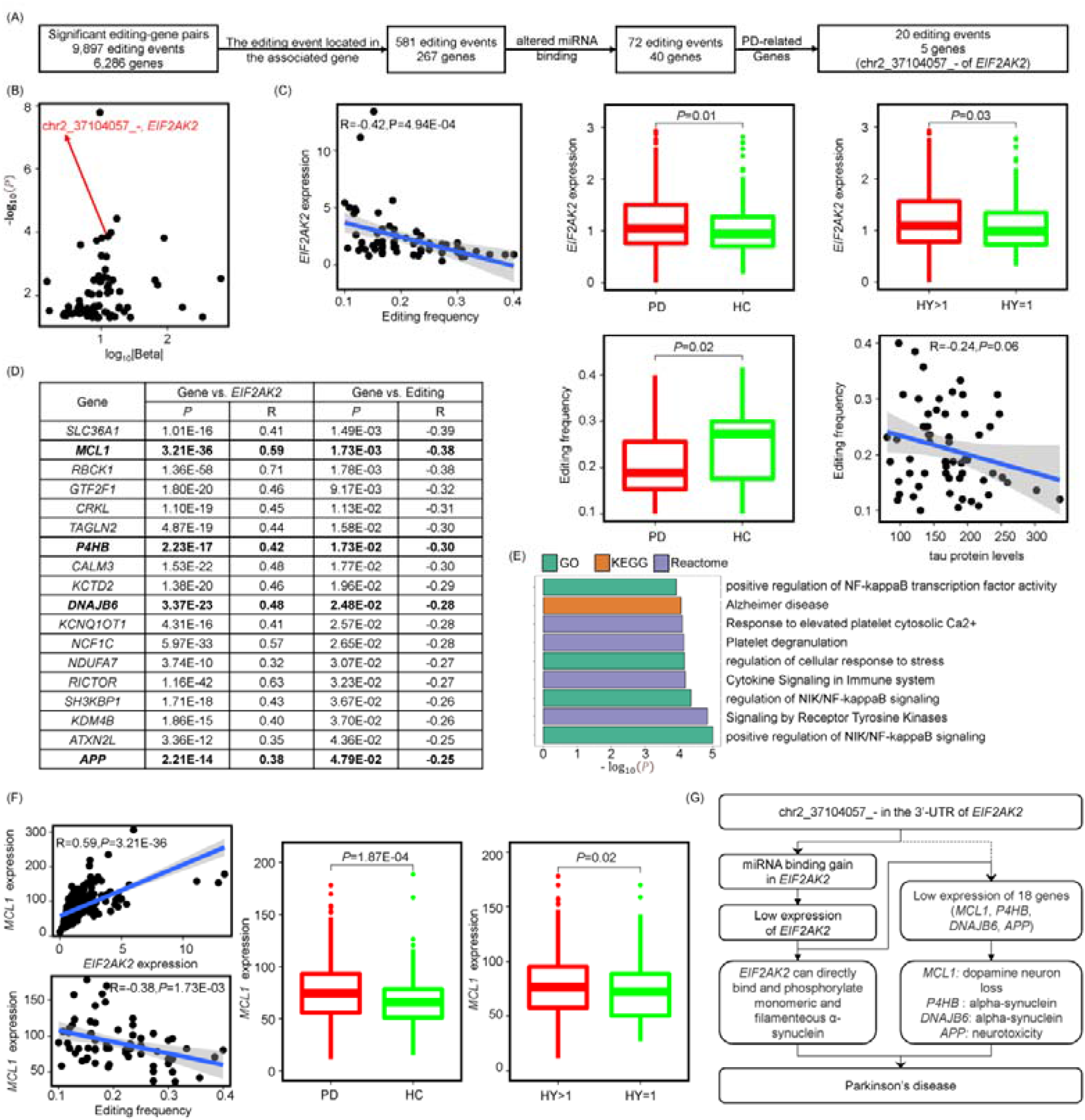
A-to-I RNA editing events may affect their host genes by interfering in miRNA regulations. (A) Analysis procedures for the underlying mechanisms of RNA editing events to be associated with their host genes. (B) The scatter plot showed the RNA editing events which may alter miRNA regulations on their host genes, highlighting one editing example in *EIF2AK2*. (C) This RNA editing event was negatively associated with *EIF2AK2* expression. Since *EIF2AK2* was highly expressed in PD samples compared to controls, and also in more sever PD samples characterized by H&Y stages, the RNA editing event may play important roles in Parkinson’s disease through its effects on this PD-related gene. It was also supported by the lower editing frequencies in PD samples and negative associations with tau protein levels. (D) Since *EIF2AK2* was an RNA binding protein, we performed correlation analyses between its potential targets and *EIF2AK2* or the RNA editing event. (E) The genes probably affected by this dys-regulated RNA binding protein were enriched in neurodegeneration related processes. (F) Of them, *MCL1* was overexpressed in PD and more sever PD samples. (G) The analyses above may uncover the potential roles of this RNA editing event in Parkinson’s disease though altered miRNA regulation.

This RNA editing event located in Chr2:37104057 position of *EIF2AK2*. It created new binding targets of *miR-3622a-3p* and *miR-3622b-3p* to probably increase the degradations of this gene. It was supported by their negative associations tested by QTL analysis (FDR = 1.39E-04, Beta = −12.60) and Pearson correlation analysis (*P* = 4.94E-04, R = −0.42) shown in **Figure 3C**. For this gene, we discovered its abnormally higher expressions in PD (*P* = 0.01) and more sever PD samples (*P* = 0.03), and its functions in promoting Parkinson’s progression through the phosphorylation of alpha-synuclein protein [13]. We may suggest this RNA editing event as a potential biomarker in Parkinson’s disease through the down-regulations of its host gene. This hypothesis was also backed by the significantly reduced frequencies of this editing event in PD samples compared to controls (*P* = 0.02), and weakly negative associations between editing frequencies and tau protein levels (*P* = 0.06, R = −0.24).

To further explore the biological functions and involved pathways of this RNA editing event, we identified 91 RNAs interacting with its host gene from StarBase [43], since this gene was proposed as an RNA binding protein [48]. Of them, 18 RNAs were demonstrated to be associated with both of this editing event and its host gene (**Figure 3D**). These genes together with the edited gene were mainly enriched in neurodegeneration related NIK/NF-KB signaling and immune processes (**Figure 3E**, **Table S8**). For example, the NF-KB plays prominent roles in dopaminergic neurodegeneration [49]. And the immune dysregulation can cause the upregulation of inflammatory cytokines to initiate a cascade of pro-inflammatory signaling that ultimately result in the neurotoxicity related to Parkinson’s disease [6]. Thus, the RNA editing event may also involve in these functions through its effects on the host gene and multiple other genes, especially on the five PD-related genes including *EIF2AK2* [13], *MCL1* [15], *P4HB* [50], *DNAJB6* [51], and *APP* [52].

Specifically, beside the host gene, *MCL1* was studied to be associated with the loss of dopamine neurons, one reason for the motor symptoms of Parkinson’s disease [15]. Moreover, in our study, we also discovered its over-expressions in PD samples (P = 1.87E-04) and that with sever conditions (P = 0.02). Its possible regulation by the RNA editing event was validated by the co-expressions of this gene and the edited gene (*P* = 3.21E-36, R = 0.59), and its negative correlations with the editing event (*P* = 1.73E-03, R = −0.38), as shown in **Figure 3F**. Based on the analyses above, we could annotate one potential mechanism that RNA editing events in miRNA targets could disturb miRNA degradation functions on their host genes and then affect the associations between these genes and others (**Figure 3G**).

### A-to-I RNA editing events may alter miRNA competition between their host genes and other genes

Beside RNA editing effects on RNA binding proteins to be associated with multiple genes, especially PD-related genes, there were also possible RNA editing events which may affect other genes though altering their miRNA competitions with the edited genes. Combing the results of QTL, Pearson correlation, and miRNA binding prediction, we identified eight RNA editing events which may alter miRNA regulations on their host genes, accompanied by their effects on 1,146 miRNA competing genes (**Figure 4A, Table S9**). Among these potentially affected genes, 127 were associated with Parkinson’s disease, involving in the key PD-related biological processes, such as necroptosis, autophagy, apoptosis, and dopaminergic synapse (**Figure 4B**, **Table S10**). These genes were possibly affected by miRNA dysregulations due to four RNA editing events including one in Chr22:35661178 position of *APOL6*.

**Figure 4.**
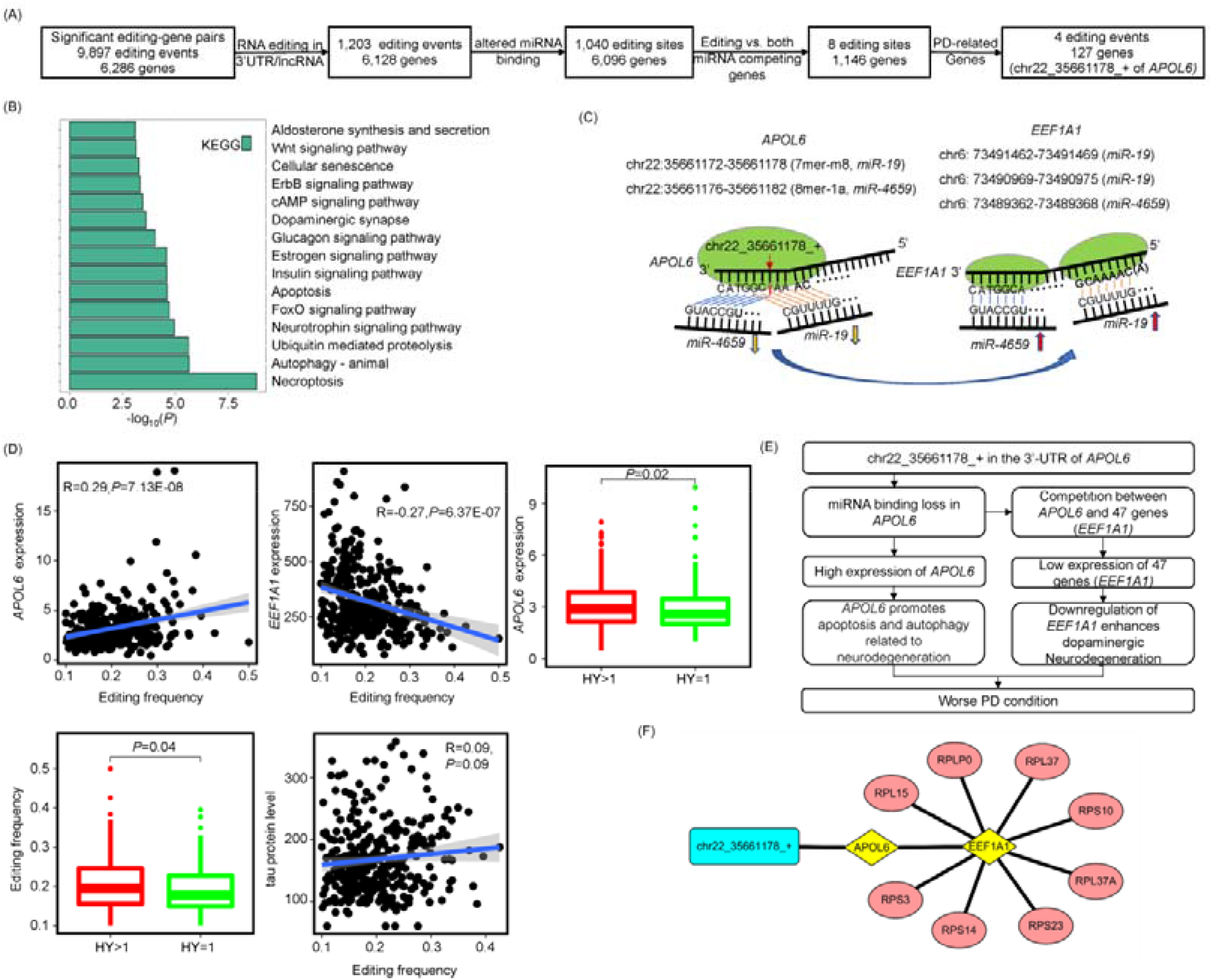
A-to-I RNA editing events may alter miRNA competitions between their host genes and other genes. (A) Analysis procedures for one underlying mechanism of RNA editing events to be associated with other genes. (B) The enrichment results of altered PD-related genes associated with the RNA editing events through miRNA competitions. (C) One RNA editing event in the 3’-UTR of *APOL6* caused the binding loss of *miR-19* and *miR-4659* to have an effect on *EEF1A1* with the binding targets of same miRNAs. (D-E) The lost miRNA binding directly led to the over-expressions of the edited gene, while indirectly caused the down-regulations of *EEF1A1*. Due to the involvements of these two genes in Parkinson’s disease, this RNA editing event may be a biomarker of the disease pathogenesis. It was supported by its higher editing frequencies in the samples with higher H&Y stages or tau protein levels. (F) This RNA editing event may have further effects on eight ribosomal proteins from their interactions with *EEF1A1*.

This RNA editing event located in the 3’-UTR of *APOL6*, causing the binding loss of *miR-19* and *miR-4659*, as shown in **Figure 4C**. The lost miRNA regulations then directly increased the expressions of *APOL6* (QTL: FDR = 1.84E-04, Beta = 8.55; Pearson: *P* = 7.13E-08, R = 0.29), while in-directly decreased the expressions of 47 genes including *EEF1A1* (QTL: FDR = 2.29E-03, Beta = −609.93; Pearson: *P* = 6.37E-07, R = −0.27) due to their competitions (**Figure 4D**). For the edited gene, we discovered its inducing roles in neurodegeneration related apoptosis process [53–55] and over-expressions in more sever PD samples (P=0.02). Moreover, the other PD-related gene competing with the edited gene, *EEF1A1*, presented its effectively pro-survival function in protective dopaminergic neurons, whose death was one characteristic of Parkinson’s disease [16]. Based on these analyses and also higher editing frequencies in more sever PD samples characterized by H&Y stages (P=0.04), and tau protein levels (*P* = 0.09, R = 0.09), we may suggest this RNA editing event as a pathogenic biomarker in Parkinson’s disease though the up-regulations of *APOL6* and down-regulations of 47 genes including *EEF1A1* (**Figure 4E**).

The *EEF1A1* gene is a translation factor responsible for the protein synthesis by delivery of aminoacyl tRNAs to the ribosome. Consistent with its functions, we also identified eight ribosomal proteins (**Figure 4F**) possibly affected by this RNA editing event though their interactions with dys-regulated *EEF1A1* [43]. Thus, the involvements of this RNA editing event in the biosynthetic and metabolic process of peptides, especially PD-related peptides, might be determined. All the analyses in this section revealed one possible mechanism for the trans-regulations of A-to-I RNA editing events on genes through the alterations of miRNA competition relationships.

### A-to-I RNA editing events may modify the seed regions to disturb miRNA regulations

It is well-known that the miRNA seed region is highly conserved that a single nucleotide change in it would dramatically alter the degradation functions of this miRNA on other genes. Thus, it is important to study the potentials of RNA editing events in miRNA seed regions to show their effects on the expressions of miRNA targets. After the analysis, we only identified one RNA editing event in Chr9:63819627 position of *miR-4477b* which dysregulated four genes (**Table S11**), including the PD-related gene of *AGO2* (**Figure 5A**).

**Figure 5.**
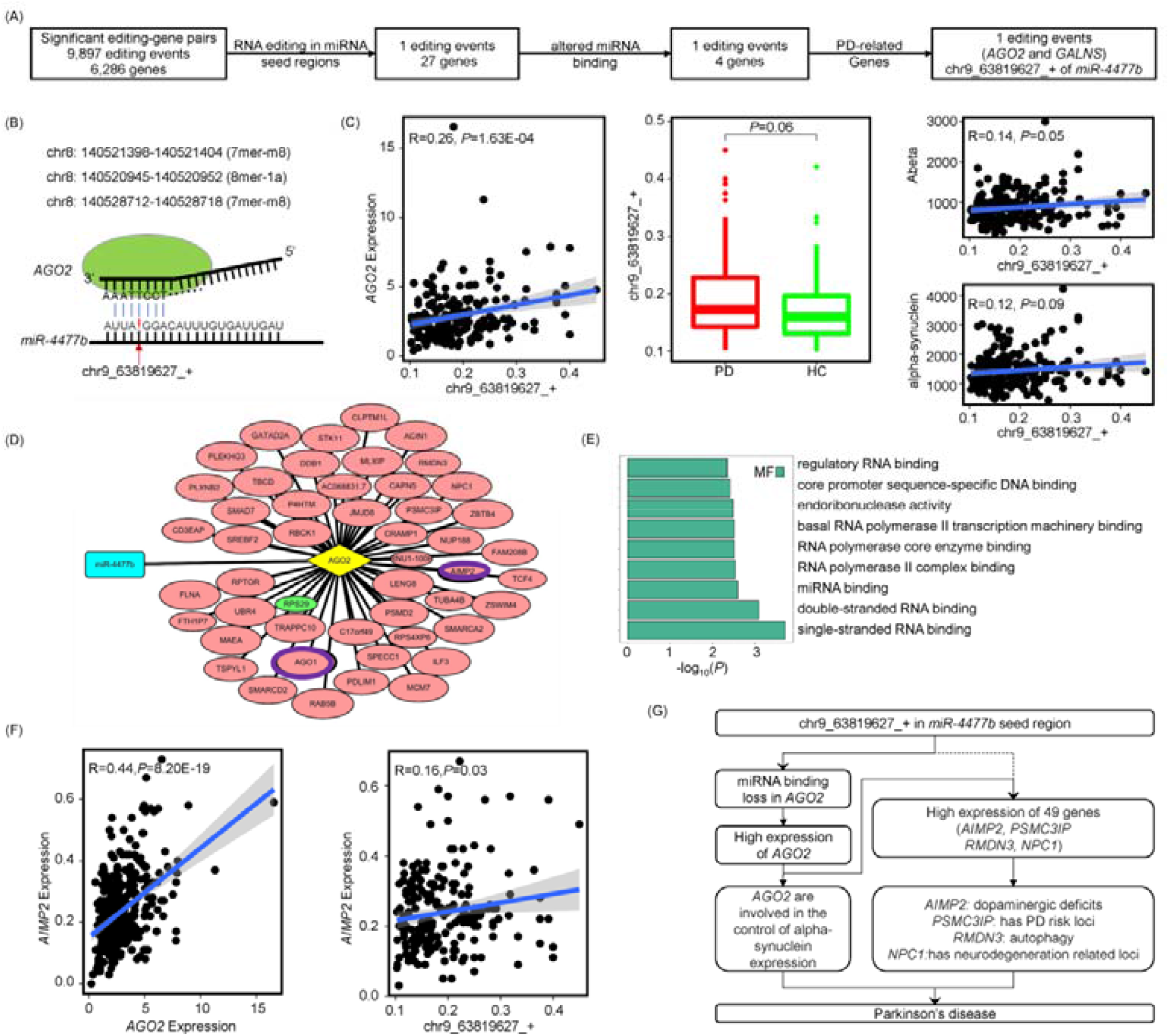
A-to-I RNA editing events may modify the seed regions to disturb miRNA regulations. (A) Analysis procedures for the underlying mechanisms of RNA editing events in miRNA seed regions to be associated with their targets, highlighting one editing example in *miR-4477b*. (B) This RNA editing event altered the miRNA seed region, thus leading to the loss of miRNA binding and regulations on *AGO2*. (C) The lost miRNA regulation caused the higher expressions of this PD-related gene. Thus, this RNA editing event may be a biomarker of Parkinson’s disease, validated by its higher frequencies in PD samples and weakly positive associations with the levels of Abeta and alpha-synuclein. (D) Since *AGO2* was an RNA binding protein, we performed correlation analyses to identify 49 potential targets associated with *AGO2* and the RNA editing event. The red and green circles described positive and negative associations respectively. (E) The discovered *AGO1* and *AGO2* genes were enriched in the miRNA and RNA binding regulations, revealing the molecular functions of *AGO2*. (F) One PD-related gene, *AIMP2*, was discovered to be associated with this RNA binding protein and RNA editing event. (G) The analyses above may uncover the potential roles of this RNA editing event in Parkinson’s pathogenesis through modifying miRNA competitions of important PD-related genes.

This RNA editing event would lead to the loss of three binding targets in *AGO2*, due to the nucleotide change from inosine to guanine in the fourth position of the miRNA seed region (**Figure 5B**). The lost miRNA regulations then caused the increased expressions of this gene (QTL: FDR = 0.02, Beta = 7.21; Pearson: *P* = 1.63E-04, R = 0.26, **Figure 5C**). Given that this gene was involved in the control of alpha-synuclein production reported in previous study [12], we may reveal the roles and pathways of this RNA editing event in Parkinson’s pathogenesis. This was also supported by the higher editing frequencies in PD samples compared to controls (*P* = 0.06) and weakly positive associations of this event with the levels of amyloid β-protein (*P* = 0.05, R=0.14) and alpha-synuclein (*P* = 0.09, R=0.12).

Furthermore, we also collected 168 RNAs interacting with this gene from StarBase, since *AGO2* was an RNA binding protein. Through the correlation analyses of these interacted genes with *AGO2* and the editing event, we identified 49 genes which may be affected by the edited miRNA and dys-regulated *AGO2* (**Figure 5D**). Among them, the argonaut family of proteins, such as *AGO1* and *AGO2*, play a role in short-interfering-RNA-mediated gene silencing (**Figure 5E**). Besides, there were also four PD-related genes probably involved in this regulation process, such as the significant associations of *AIMP2* with *AGO2* and the editing event (**Figure 5F**). They showed their mutation associations with neurodegenerative diseases [56–57], activation roles in age dependent dopaminergic neuronal loss [17], and regulatory function of PD-related autophagy [58]. All the above analyses gave the possible evidence that RNA editing events in miRNA seed regions may affect their degradation functions on multiple genes to be involved in Parkinson’s disease (**Figure 5G**).

## DISCUSSION

The A-to-I RNA editing events in Parkinson’s disease have attracted the interests of researchers in recent years [20,59]. For now, these studies mainly focused on the differential RNA editing events in coding regions and miRNAs. The further functional roles of these RNA editing events and also the events in lncRNAs or 3’-UTRs of mRNAs needed to be addressed. Here, this study analyzed 372 PD patients for the identification of 9,897 A-to-I RNA editing events associated with 6,286 genes. Of them, 820 genes were proposed previously to be linked with Parkinson’s disease. Since most of all these associations belonged to the trans-regulatory group (**Figure 2**), it was urgent and necessary to understand the underlying mechanisms of the PD-related genes to play their roles in Parkinson’s pathogenesis. Here, we explored that from the view of disturbed miRNA regulations on genes due to A-to-I RNA editing events. Specifically, we identified 72 RNA editing events probably interfering in miRNA regulations on their host genes (**Table S7**), eight RNA editing events possibly altering miRNA competitions between their host genes and 1,146 other genes (**Table S9**), and one RNA editing event modifying miRNA seed regions to potentially disturb its regulations on four genes (**Table S11**). These analyses provided one kind of mechanisms for the roles of 25 A-to-I RNA editing events to alter miRNA regulations on 133 PD-related genes.

During the analyses, to increase the reliability of the discovery, we performed both of QTL and Pearson correlation tests for the identification of the associations between A-to-I RNA editing events and genes. The associations were determined only when they all passed the statistical significance thresholds of the two tests. As for the miRNA related mechanism underlying these associations, we used two miRNA binding prediction tools, TargetScan and miRanda, to identify the effects of RNA editing events on genes by altering the miRNA regulations. The gain of miRNA binding target was recognized only when the interactions occurred in RNA-edited sequences but not in wild-type sequences supported by both tools, similarly for the loss of miRNA binding target. These processes all enhanced the accuracy of the identified RNA editing events to probably interfere in miRNA regulations on genes in Parkinson’s pathogenesis.

Moreover, we also checked RNA editing effects on genes in a large dataset of Alzheimer’s disease (AD). Among the three examples introduced in detail, two RNA editing events in the 3’-UTR of *EIF2AK2* and *miR-4477b* seed region also presented their alteration roles of miRNA degradation functions on corresponding genes in the brain regions of AD patients (**Figure S2**). It revealed the reliability of this mechanism that A-to-I RNA editing events may alter miRNA regulations on genes. On the other hand, it also showed the consistently regulatory functions of A-to-I RNA editing events in neurodegenerative diseases. Combined the functions of RNA editing events in Alzheimer’s disease [21], this study will expand the knowledge for the RNA editing effects on neurodegeneration related genes.

The study of RNA editing effects on key genes pointed out the RNA editing biomarkers in neurodegenerative diseases. To improve the understanding of these biomarkers, it is better to annotate their potential up-regulators. Specifically, for the three editing examples introduced in detail, we discovered the main editing enzyme and genetic variants [60] which may affect these A-to-I RNA editing events (**Figure S3**). The analyses could link the DNA mutations or *ADAR*, A-to-I RNA editing events, miRNA regulations, and gene expressions together, to explain the potentially regulatory processes in neurodegenerative pathogenesis and be helpful for the treatment of the diseases.

Overall, this study explored partial involved biological processes of individual A-to-I RNA editing event in Parkinson’s disease. Given the complex molecular mechanisms of RNA editing events, we plan to further study their effects in Parkinson’s disease from the following two aspects. First, we will build an editing-expression network for their potential interactions, since we discovered the possibility of multiple RNA editing events interfering in miRNA regulations on multiple genes shown in **Figure S4**. Second, we will further explore another kind of mechanisms that RNA editing events may disturb the regulations of RNA binding proteins on genes. Initially, we have identified 140 RNA editing events in RNA binding proteins and 4,182 RNA editing events in their targets from StarBase [43]. Interesting, one RNA editing example introduced here seemed also involved in *HNRNPA1* regulations on *APOL6* and *EEF1A1* (**Figure S5**). These two future studies will improve the understanding of RNA editing functions in Parkinson’s pathogenesis.

## Acknowledgments

This work was supported by the National Natural Science Foundation of China (Grant No. 62002270), the Fundamental Research Funds for the Central Universities, the Natural Science Foundation of Shaanxi Province of China (Grant No. 2020JQ-332), the China Postdoctoral Science Foundation (Grant No. 2018M643583), National Key R&D Program of China (Grant No. 2017YFA0205202), and partially funded by the National Natural Science Foundation of China (Grant No. 61672422). The funders had no role in study design, data collection and analysis, decision to publish or preparation of the manuscript.

