## Supplementary Figure 1-5 for "The Potential Regulation of A-to-I RNA editing on Genes in Parkinson’s Disease"

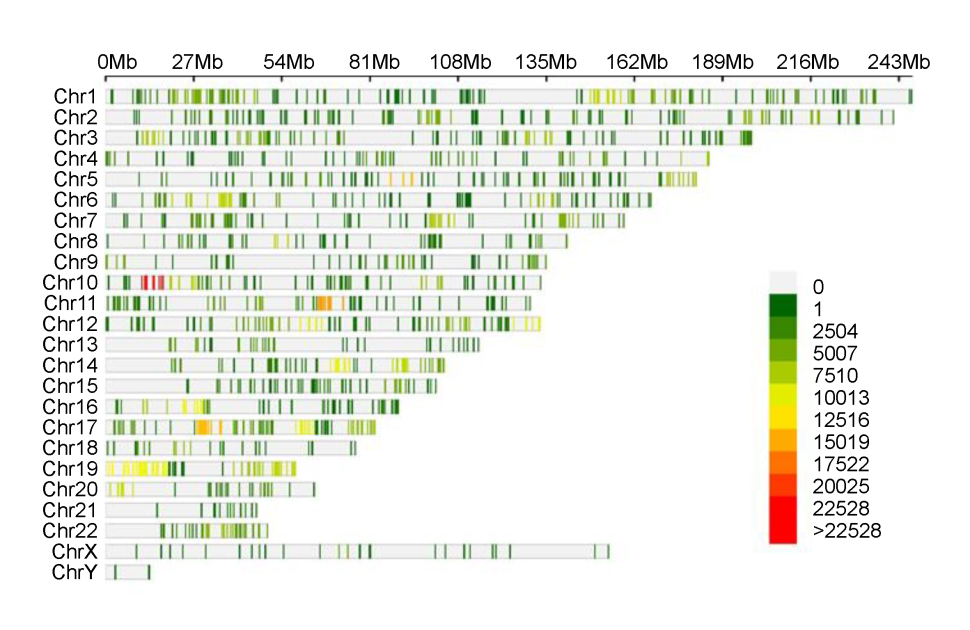
**Figure S1.** **The density of RNA editing events associated with genes in chromosomes.**

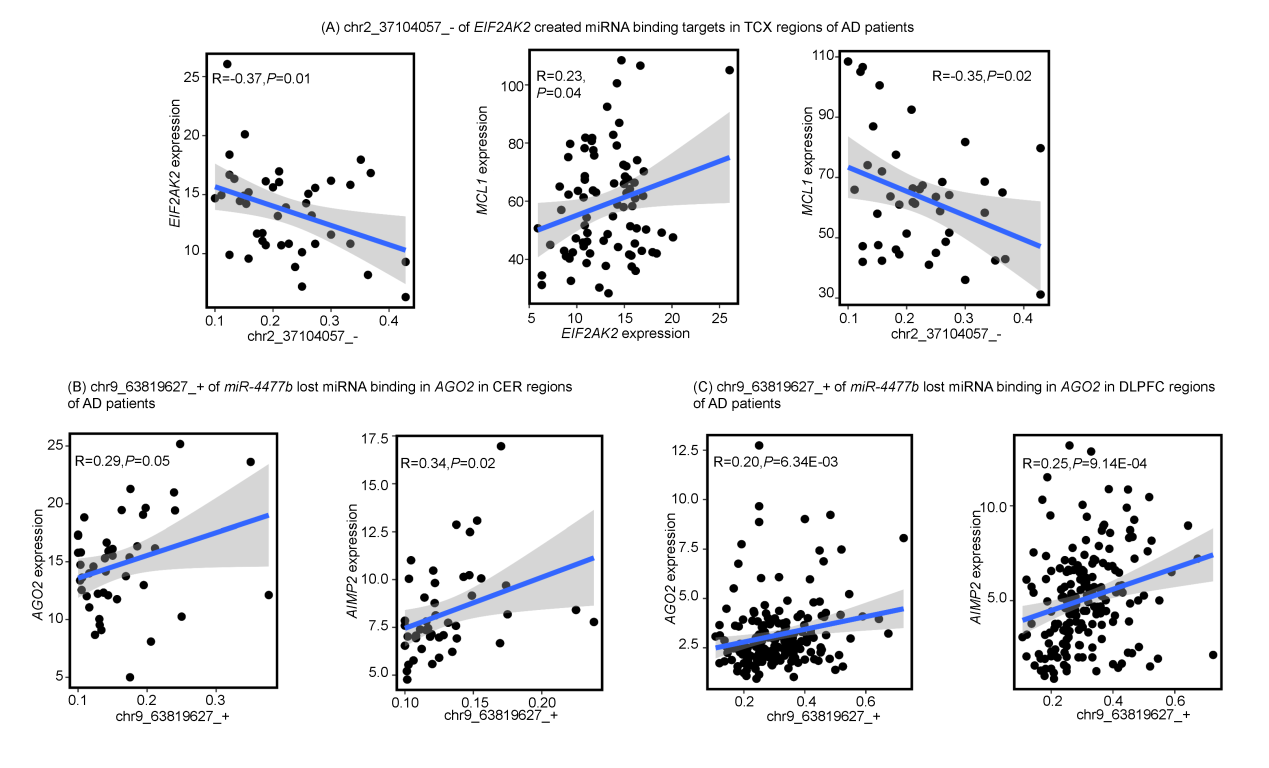
**Figure S2. The functions of RNA editing events in Alzheimer’s disease (AD)**. (A) In temporal cortex (TCX) regions of AD patients, the RNA editing event (chr2_37104057_-) in *EIF2AK2* created miRNA binding targets, leading to the reduced expressions of its host gene. Due to the interactions of *MCL1* with its host gene, this RNA editing event may also have effect on *MCL1*. (B-C) In cerebellum (CER) and dorsolateral prefrontal cortex (DLPFC) regions of AD patients, the RNA editing event (chr9_63819627_+) in the seed region of *miR-4477b* may dys-regulate *AGO2* and *AIMP2* genes.

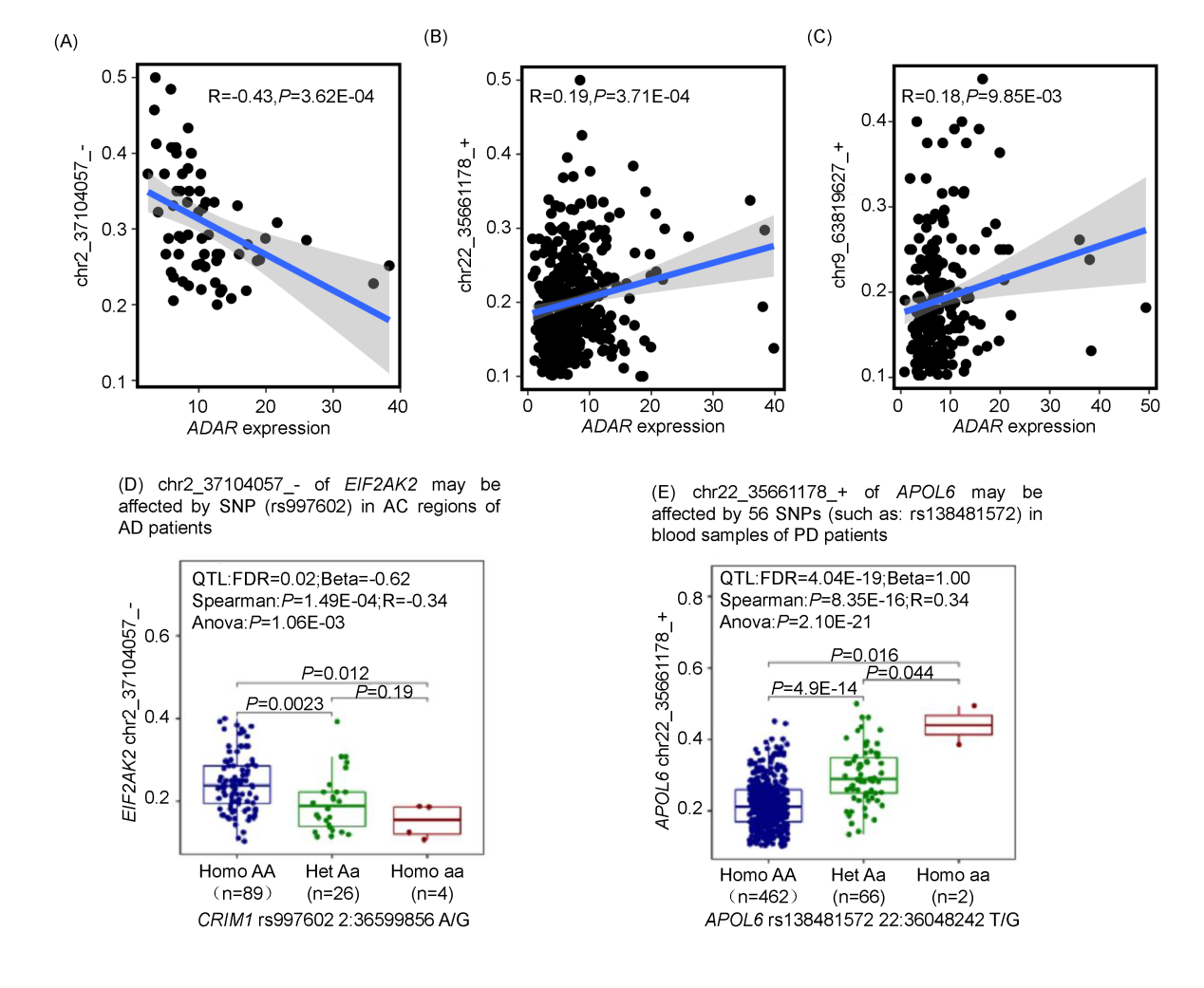
**Figure S3. The potential up-regulators of A-to-I RNA editing events**. (A-C) The correlations of *ADAR* enzyme with the three RNA editing events. (D-E) The effects of genetic variants on the two RNA editing events.

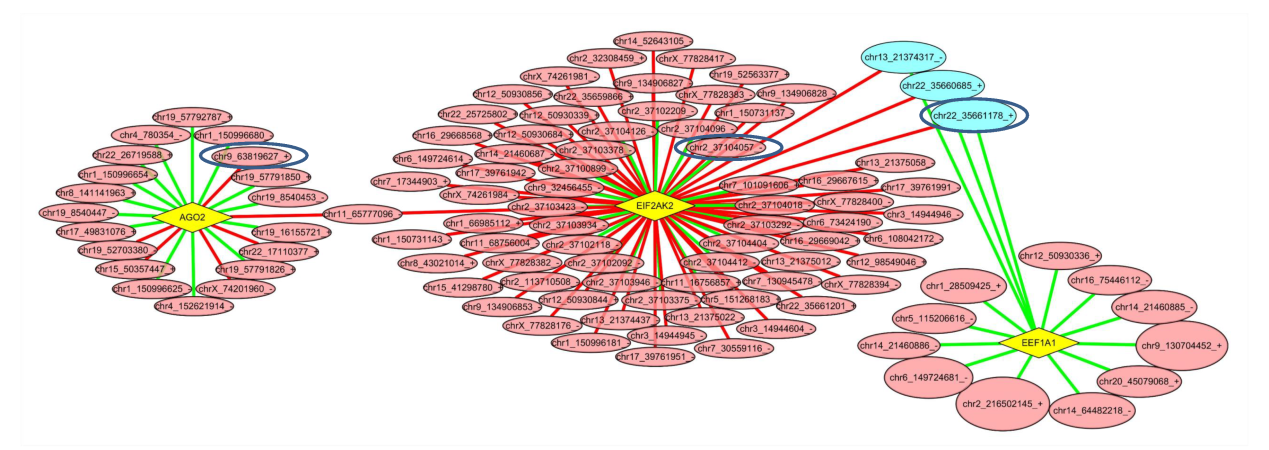
**Figure S4. The effects of multiple RNA editing events on the three genes**. These RNA editing events were all predicted to interfere in miRNA regulations. The red and green lines showed the probably positive or negative effects of RNA editing events on genes. The three RNA editing events introduced in this study were highlighted by blue circles.

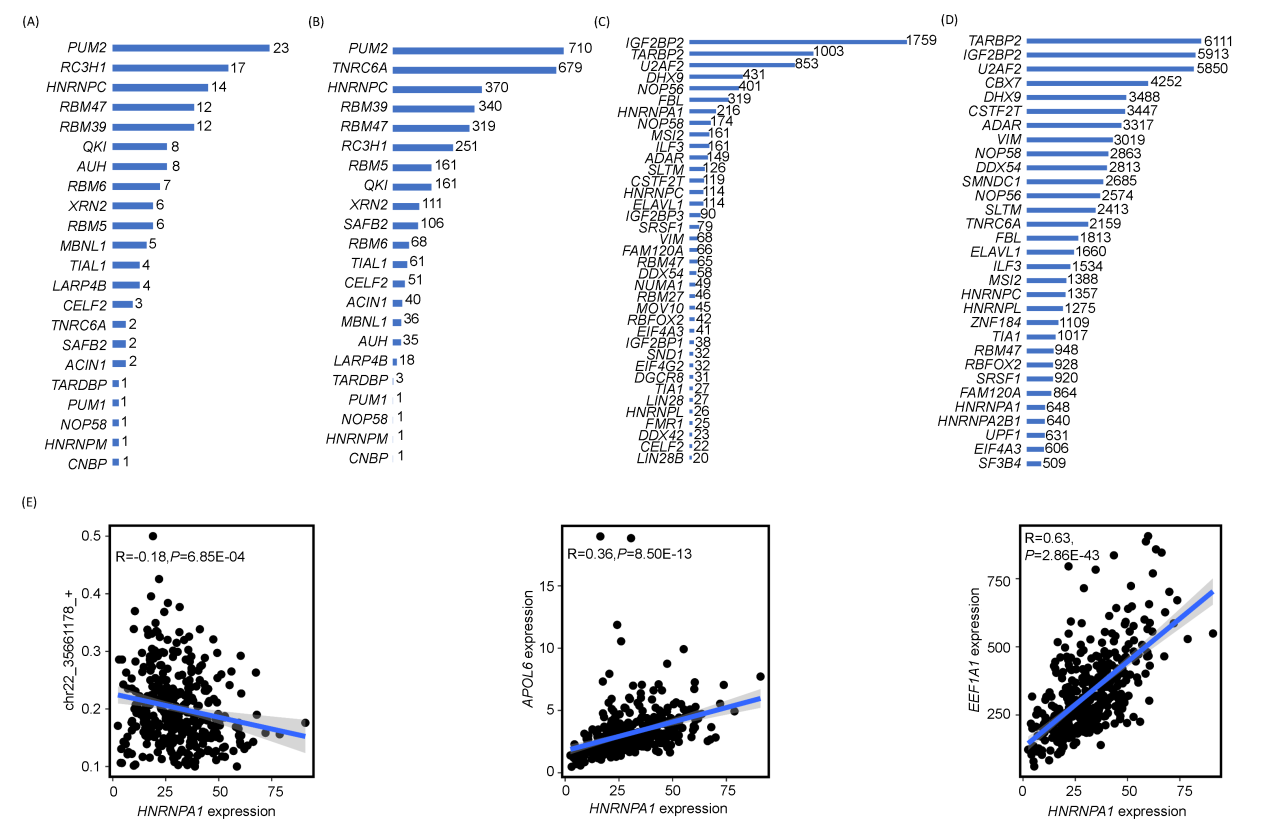
**Figure S5. The potential involvements of RNA editing events in the regulations of RNA binding proteins (RBP) on genes**. (A) The number of RNA editing events in RBPs. (B) The number of genes associated with the RNA editing events in RBPs. (C) The number of RNA editing events in RBP targets. (D) The number of genes associated with the RNA editing events in RBP targets. (E) The correlations of an RNA binding protein, *HNRNPA1*, with the editing event, *APOL6*, and *EEF1A1*. This RNA editing event located in the binding region of *HNRNPA1*, which also probably regulated the two genes.
